# Characterization of ethylene-inducible pedicel-fruit abscission zone formation in non-climacteric sweet cherry (*Prunus avium* L.)

**DOI:** 10.1101/2020.09.05.284497

**Authors:** Seanna Hewitt, Benjamin Kilian, Tyson Koepke, Jonathan Abarca, Matthew Whiting, Amit Dhingra

## Abstract

Harvesting of sweet cherry (*Prunus avium* L.) fruit is a labor-intensive process. Mechanical harvesting of sweet cherry fruit is feasible; however, it is dependent on the formation of an abscission zone at the fruit-pedicel junction. The natural propensity for pedicel-fruit abscission zone (PFAZ) formation varies by cultivar, and the general molecular basis for PFAZ formation is not well characterized. In this study, ethylene-inducible change in pedicel fruit retention force (PFRF) was recorded in a developmental time course with a concomitant analysis of the PFAZ transcriptome from three sweet cherry cultivars. In ‘Skeena’, mean PFRF for both control and treatment fruit dropped below the 0.40kg-force (3.92N) threshold for mechanical harvesting and indicating the formation of a discrete PFAZ. In ‘Bing’, mean PFRF for both control and treatment groups decreased over time. However, a mean PFRF conducive to mechanical harvesting was achieved only in the ethylene-treated fruit. While in ‘Chelan’ the mean PFRF of the control and treatment groups did not meet the threshold required for efficient mechanical harvesting. Transcriptome analysis of the PFAZ followed by the functional annotation, differential expression analysis, and gene ontology (GO) enrichment analyses of the data facilitated the identification of phytohormone-responsive and abscission-related transcripts as well as processes that exhibited differential expression and enrichment in a cultivar-dependent manner over the developmental time-course. Additionally, read alignment-based variant calling revealed several short variants in differentially expressed genes, associated with enriched gene ontologies and associated metabolic processes, lending potential insight into the genetic basis for different abscission responses between the cultivars. These results provide genetic targets for induction or inhibition of PFAZ formation, depending on the desire to harvest the fruit with or without the stem attached. Understanding the genetic mechanisms underlying the development of the PFAZ will inform future cultivar development while laying a foundation for mechanized sweet cherry harvest.

## Introduction

Sweet cherry (*Prunus avium* L.), a member of the Rosaceae family, is a commercially important tree fruit crop throughout the world, with approximately 2.3 million tons produced annually (Axe and Bush, 2017; Blando and Oomah, 2019). In addition to its worldwide economic value, production, and consumption of sweet cherries has increased in recent years as consumers have become aware of their nutritional benefits (Blando and Oomah, 2019). The industry is evolving to address the growing market, availability of labor during harvest, and has begun adapting harvesting strategies to meet new consumer demand for stemless cherries (Kappel et al., 2012; Quero-García et al., 2017). Traditional harvesting methods focus on separating the fruit from the tree at the pedicel-peduncle junction, leaving the pedicel (stem) attached to the fruit (Wittenbach and Bukovac, 1974). Mechanical harvesting, on the other hand, is best achieved when the fruit abscises easily at the fruit-pedicel junction (Quero-García et al., 2017). Increasing labor costs associated with traditional hand harvesting, in addition to the growing demand for stemless fruit, has made adoption of mechanical harvesting strategies attractive if they can be made uniform across cultivars (Zhao et al., 2013). However, the harvesting of sweet cherries has presented a unique set of challenges. Unlike sour cherries (*Prunus cerasus*), which develop an anatomically and histologically distinct fruit-pedicel abscission zone (PFAZ) and separate with ease, sweet cherry cultivars display phenotypic differences in PFAZ formation and consequent ease of fruit separation—some cultivars require excessive force to separate fruit at the PFAZ, which tends to compromise fruit quality and integrity (Stösser et al., 1969; Zhao et al., 2013).

While sweet cherry and peach belong to the same sub-family, the former bears non-climacteric fruits that do not produce ethylene autocatalytically during ripening and senescence. This is most likely due to the presence of several stop codon mutations in the ethylene biosynthesis and perception genes in sweet cherry (Koepke et al., 2013). However, sweet cherry fruit from some cultivars exhibit a novel developmental response to exogenous ethylene application (Hiwasa-Tanase and Ezura, 2014). Exogenous ethylene application can induce or enhance PFAZ formation, loosening the fruit, and facilitating efficient mechanical harvesting (Wittenbach and Bukovac, 1974; Smith and Whiting, 2010).

Pedicel-fruit retention force (PFRF) is used as a direct measure of PFAZ formation and serves as a metric to determine the mechanical harvestability of the fruit. An average PFRF value of 0.40kg-force (3.92N) is considered the threshold for mechanical harvestability, though this will depend on the actuation method of the harvester (Bukovac and MJ, 1979; Smith and Whiting, 2010; Zhao et al., 2013). In sweet cherry, a reduction in PFRF can be induced through the application of ethephon (2-chloroethylephosphonic acid), a commercially available plant growth regulator that is rapidly metabolized to ethylene (Smith and Whiting, 2010; Zhao et al., 2013). In both sweet and sour cherry, ethephon has been demonstrated to be a viable option for reducing the threshold for mechanical harvest without negatively impacting fruit quality, when applied in appropriate concentrations; however high rates of ethephon have been associated with reduction in fruit quality (e.g. gummosis, defoliation, lenticel enlargement) and, thus, cultivars with naturally low PFRF are desired (Looney and McMechan, 1970; Bukovac and MJ, 1979; Smith and Whiting, 2010; Zhao et al., 2013). The natural PFRF of sweet cherry varies by cultivar, with some varieties, like ‘Skeena’, exhibiting an auto-abscising phenotype and requiring no exogenous ethylene to induce abscission. Representing an intermediate phenotype, ‘Bing’ can be induced to abscise when ethephon is applied approximately 14 days before harvest (Smith and Whiting, 2010; Zhao et al., 2013). ‘Chelan’, on the contrary, does not abscise naturally or in the presence of ethephon (Smith and Whiting, 2010).

Sweet cherry phenotyping studies have shown that PFRF values remain consistent for these cultivars across multiple years, indicating that the abscission phenotypes are genetically stable and can perhaps be manipulated at the genetic level (Zhao et al., 2013). Furthermore, these findings suggest that standardization of PFRF for mechanical harvesting across cultivars is possible if the ideal ethephon or other treatment regimens are determined for individual sweet cherry cultivars. Despite the extensive physiological characterization of PFAZ integrity across sweet cherry varieties, the underlying molecular basis for PFAZ structural differences has not previously been elucidated.

As understood from studies in model plant systems such as Arabidopsis and tomato, abscission at the fruit-pedicel junction entails a series of hallmark structural changes: the middle lamella is dissolved by hydrolytic enzymes, such as polygalacturonase and cellulase (Taylor et al., 1991); cell walls in the separation layer thicken, and cell wall components become hydrated as a result (Huberman et al., 1983); primary cell walls break down as abscission progresses, resulting in the formation of large intercellular cavities (Tabuchi et al., 2001; Grover, 2012); and lignin deposits accumulate proximally to the abscission zone, forming part of a peridermal boundary layer that will serve to protect the pedicel scar following fruit separation (Sexton and Roberts, 1982; Tabuchi et al., 2001; Merelo et al., 2017; Meir et al., 2019). This process is thought to operate in a similar manner in fruit crops including apple, peach, and olive although species and cultivar-specific differences, particularly with regards to chemical induction of abscission in these crops, are not yet well understood (Botton et al., 2011; Zhu et al., 2011; Ali et al., 2012; Gil-Amado and Gomez-Jimenez, 2013; Goldental-Cohen et al., 2017).

According to the current model of abscission in plants, the genetic events underlying cellular structure modification at the PFAZ center around an interplay between ethylene and auxin-associated pathways. Binding of ethylene to corresponding receptors (ETRs) in PFAZ cells initiates signal transduction pathways leading to the activation of numerous, ethylene-responsive transcription factors (ERFs), which elicit different modulatory roles. The ethylene response and signaling network ultimately results in the initiation of cell death, reduction in cell wall adhesion, and separation of the fruit from the pedicel (Roberts et al., 2002; Meir et al., 2019). Working antagonistically to ethylene is the phytohormone auxin. Auxin’s biologically active form, free indole-3-acetic acid (IAA), decreases the sensitivity of plant organs to ethylene. The genetic and metabolic factors governing auxin homeostasis ensure that an appropriate balance is maintained between free IAA, IAA-conjugates, and auxin degradation during different developmental stages (Meir et al., 2010, 2015). A presence of higher levels of free IAA corresponds to inhibited or delayed ethylene-dependent developmental responses like abscission zone formation (Else et al., 2004). Decreasing polar auxin transport across the abscission zone in sweet cherries by girdling methods results in increased fruit abscission (Blanusa et al., 2005). Additionally, in grape, the application of inhibitors of auxin transport has been shown to promote increased abscission (Kühn et al., 2016).

While there is evidence implicating the involvement of numerous ethylene-associated transcription factors and auxin-associated genes in the modulation of PFAZ formation, species-specific modes of action have yet to be resolved (Roberts et al., 2002). Furthermore, mechanisms involving transmission of hormonal and other signals upstream of AZ formation, as well as specifics of the enzyme-driven remodeling and redeposition of cell wall constituents in sweet cherry are in need of further elucidation. The stimulation of PFAZ formation in response to exogenous ethylene application in the non-climacteric fruit sweet cherry represents a unique biological system to elucidate the process of inducible abscission. An improved understanding of the interplay between hormone response, signaling, and activation of abscission-associated genes and pathways in sweet cherry will facilitate improved strategies for planned induction or inhibition of PFAZ formation.

To elucidate the molecular bases for differences in abscission phenotypes among sweet cherry cultivars, and to correlate this information with fruit development, time course physiological measurements of the PFRF along with concomitant transcriptome analysis of the PFAZ tissue from ethylene-treated and control ‘Bing’, ‘Skeena’, and ‘Chelan’ were conducted. The hypothesis that cultivar-specific gene expression differences in ethylene- and auxin-responsive pathways are directly correlated to the differences in abscission phenotypes was evaluated. Additionally, this work aimed to reveal other genes and enriched gene ontologies associated with key metabolic processes involved in pedicel-fruit abscission in sweet cherry. The results of this study provide potential genetic targets for PFAZ formation in sweet cherry, which are expected to inform strategies for improving PFAZ phenotypes conducive to different harvesting approaches.

## Methods

### Plant Material

The sweet cherry trees used in this study are located at Washington State University’s Roza Farm, 10 km north of Prosser, Washington, USA (46.2°N, 119.7°). Trees were irrigated weekly from bloom to leaf senescence with low-volume, under-tree, micro-sprinklers, and grown using standard orchard management practices. Trees had an in-row spacing of 2.44 m (8 ft) and between row spacing of 4.27 m (14 ft). Rows were planted in a north-south orientation and trained to a Y-trellis architecture.

### Ethephon application

Ethephon (formula 240 g/l [2lbs/gal]) was applied via air-blast sprayer at 3.5 L ha−1 (3 pt A−1) with a 1,871 L ha−1 (200 g A−1) spray volume (Smith and Whiting, 2010). Ethephon applications and measurements were conducted in three different years (2010, 2013, and 2014). Each replication was performed in the same orchard block, using distinct trees within the block. Treatment application was done early in the morning (between 0600 and 0800 hours) to reduce the effects of ethylene evolution from warm temperatures and wind, as previously described (Smith and Whiting, 2010).

Optimal ethephon application time for ‘Bing’ had been established previously as 14 days before harvest (DBH) (Smith and Whiting, 2010), or 80% fruit maturation. Because ‘Bing’, ‘Chelan’, and ‘Skeena’ have different timelines for the maturation of fruit after bloom, ethephon was applied at ca. 80% maturation for each of the cultivars in the 2014 growing season. This percentage coincided with 14 DBH for ‘Bing’, 12 DBH for ‘Chelan’ and 16 DBH for ‘Skeena’ (Supplementary File 1). Information regarding ethephon treatment, and PFRF results for 2010 and 2013 can be found in Supplementary File 2.

### PFRF measurements, color evaluation, and abscission zone sampling

In all three years, sampling and measurements were conducted at the following time points for each sweet cherry cultivar: (1) immediately prior to application of ethephon or H_2_O; (2) 6 hours after the application of ethephon or H_2_O, and (3) at harvest.

At each sampling time, ten fruit were randomly selected for analysis from each of four trees/cultivar/treatment. PFRF was measured using a modified digital force gauge (Imada). Each fruit was manually categorized by exocarp color based on 1–7 scale developed specifically for sweet cherry by CTIFL (Centre technique interprofessionnel des fruit et legumes, France) (Supplementary File 3) (Zhang and Whiting, 2011). In addition to the collection of PFRF values, the abscission zones of 10 fruit, representing ten biological replicates, from each cultivar/treatment were harvested from corresponding trees at each time point per the following steps: 1.) Using a new razor blade, the fruit was first cut approximately 0.5 cm below the pedicel, leaving the pedicel and a thin disc of fruit/skin attached, 2.) two sets of parallel cuts were made downward on the cherry fruit disc on either side of the stem, effectively making a cubic piece of fruit 3mm x 3mm x 3mm attached to the pedicel, 3.) the pedicel was cut off directly above the fruit and the cube of fruit tissue consisting the abscission zone and some fruit and pedicel tissue was placed in a 15ml falcon tube and flash frozen for subsequent processing (Supplementary File 4).

### Total RNA Extractio**n**

Excised sweet cherry abscission zone tissue derived from ten fruits representing ten biological replicates / cultivar / time point was pooled, pulverized and homogenized into a single sample using a SPEX SamplePrep® FreezerMill 6870 (Metuchen, NJ USA) and transferred to storage at -80°C. Total RNA was extracted using an acid guanidinium thiocyanate phenol chloroform extraction method similar to that previously described (Chomczynski and Sacchi, 1987). Briefly, 1mL of 0.8M guanidinium thiocyanate, 0.4M ammonium thiocyanate, 0.1M sodium acetate pH 5.0, 5% w/v glycerol, and 38% v/v water saturated phenol were added to approximately 100 mg powdered tissue, shaken to evenly mix the sample and incubated at room temperature (RT) for 5 minutes. 200μL chloroform was added and shaken vigorously until the entire sample became homogenously cloudy and then was incubated (RT, 3 minutes). Samples were centrifuged at 17,000 x g at 4°C for 15 minutes, and the aqueous upper phase was transferred to a sterile 1.5mL microcentrifuge tube. To this, 600μl 2-propanol was added, inverted 5-6 times, and incubated at RT for 10 minutes. Samples were centrifuged 17,000 x g at 4°C for 10 minutes, and the supernatant decanted. 1 mL 75% DEPC ethanol was added to the pellet, vortexed for 10 seconds, and centrifuged 9,500 x g at 4°C for 5 minutes. Pellets were suspended in RNase free water and incubated at 37°C with RNase free *DNaseI* for 30 minutes, which was subsequently inactivated (65°C, 10 minutes).

Extracted RNA was quantified, and its quality was checked using the Bio-Rad (Hercules, CA) Experion system using the Experion RNA High Sensitivity Analysis kit or the Agilent (Santa Clara, CA) 2100 Bioanalyzer system using the RNA NanoChip and Plant RNA Nano Assay Class.

### RNA sequencing and assembly

RNA samples that passed the quality threshold of RIN 8.0 were sequenced at the Michigan State University Genomics Service Center for library preparation and sequencing. The Illumina Hi Seq 2000 sequencing platform (San Diego, CA) was used to sequence 2×100 PE reads from the cDNA libraries generated from the above RNA extractions, representing a single sample derived from 10 biological replicates of each cultivar, treatment, and time point. cDNA and final sequencing library molecules were generated with Illumina’s TruSeq RNA Sample Preparation v2 kit (San Diego, CA) and instructions with minor modifications. Modifications to the published protocol include a decrease in the mRNA fragmentation incubation time from 8 minutes to 30 seconds to create the final library of proper molecule size range. Additionally, Aline Biosciences’ (Woburn, MA) DNA SizeSelector-I bead-based size selection system was utilized to target final library molecules for mean size of 450 base pairs. All libraries were quantified on a Life Technologies (Carlsbad, CA) Qubit Fluorometer and qualified on an Agilent (Santa Clara, CA) 2100 Bioanalyzer.

Read preprocessing and assembly were conducted in CLC Genomics Workbench (8.5.1). Briefly, RNAseq read datasets were processed with the CLC Create Sequencing QC Report tool to assess read quality. The CLC Trim Sequence process was used to trim the first 16 bases due to GC ratio variability and for a Phred score of 30. All read datasets were trimmed of ambiguous bases. Illumina reads were then processed through the CLC Merge Overlapping Pairs tool, and all reads were *de novo* assembled to produce contiguous sequences (contigs). A single master assembly was generated from the combined read data from ‘Bing’, ‘Chelan’, and ‘Skeena’ cultivars at each time point (Supplementary File 5). Assembled contigs passed the filter criteria of >200 base length combined with >2x average read coverage. The cultivar-specific, non-trimmed read sets were mapped back to the master assembly to generate individual read mappings for each cultivar, treatment, and time point. Read counts were normalized for each mapping group using the Reads Per Kilobase per Million reads (RPKM) method (Mortazavi et al. 2008). All the sequence data is available from NCBI Short Read Archive – Bing accession: SRX2210365; Chelan accession: SRX2210366; Skeena accession: SRX2210367 submitted under BioProject PRJNA329134.

### Differential expression analysis

A two-pronged approach, using Kal’s Z-test and the NOIseq-sim R package, was employed to identify contigs with highest likelihood of being differentially expressed (Kal et al., 1999; Tarazona et al., 2013). Only contigs that passed established threshold filtering for both methods were considered for further analysis.

Genes with highest probability of being differentially expressed were first identified using the NOISeq-sim package in OmicsBox, which is designed to infer probability of differential expression by modeling biological replications in silico for RNAseq experiments (Tarazona et al., 2011, 2013). This approach has been successfully employed for differential expression analysis in other crops, including peach and rice (De La Fuente et al., 2015; Altúzar-Molina et al., 2020). Default parameters were used to simulate five, in silico replications with a simulated replicate size of 0.2 and a set variability of 0.02 in each replication.

Next, Kal’s Z-tests were performed in CLC Genomics Workbench 8.5.1 to add another level of stringency to the identification of differentially expressed genes. A paired experiment comparing the read count values for ethephon treatment to control values corresponding to each cultivar was performed.

The final sequence selection was reduced to 3,190 contigs, which were considered to have a high probability of being differentially expressed based on conformity to all of the following criteria for at least one treatment/time: (1) |log2FC|>1, (2) NOIseq probability of DE > 0.8, (3) Kal’s Z-test FDR corrected p-value <0.05 (Supplementary File 6).

### Functional annotation and GO enrichment analysis

Assembled sweet cherry contigs were annotated using the Blast2GO feature in OmicsBox (version 1.2.4). Contigs were blasted for greatest sequence homology against the NCBI Viridiplantae database and subsequently assigned to their corresponding gene ontology (GO) terms as described previously (Götz et al., 2008; Hewitt et al., 2020a, 2020b).

GO enrichment analysis using Fisher’s Exact Test was also conducted in OmicsBox to identify cellular components, molecular functions, and biological processes that were over or under-represented in the ethephon treated fruit at harvest in comparison with the control fruit (Götz et al., 2008). Based on the differential expression analysis, for each sweet cherry cultivar, lists representing transcripts with NOIseq probability>0.8, Kal’s Z-test FDR corrected p-value<0.05, and |logFC|>1 at the harvest time point were produced. These lists served as the treatment datasets for enrichment analyses, and the master annotated transcriptome was used as the reference dataset (Supplementary File 7). The FDR corrected p-value cutoff for enrichment was set to 0.05. Following separate enrichment analyses for each cultivar, enriched GO terms that were shared between cultivars or unique to a single cultivar were identified using the Venn Diagram application in OmicsBox.

### RT-qPCR Validation

Targets for RT-qPCR validation were selected from a list of genes known to be involved in ethylene response and cell wall breakdown (Supplementary File 8). Primers were designed based on the near full-length transcript sequences to amplify an approximately 100-150 bp region in the 3’ region of target transcripts. A bacterial luciferase gene was used as a spiked reference, with 50 ng added per reaction.

Library preparation, target amplification, and expression analysis were conducted in accordance with previously published methods, with minor modifications (Hendrickson et al. 2019). VILO cDNA synthesis kit (Invitrogen™) was used to generate three biological replicates of cDNA from three independent RNA samples isolated for each time point per manufacturer’s instructions. Replicate cDNAs were then pooled into a single sample (50 ng/ul). At least 4-6 technical replicate RT-qPCR reactions were performed using iTAQ with ROX and SYBR (BioRad), and 20μL reactions were prepared as per the manufacturer recommendations. A total of 2μL of cDNA diluted to 50ng/μL RNA equivalents was used per reaction with 5μL H_2_O, 2μL of each primer (10μM), and 10μL of iTAQ SYBR® Green Supermix with ROX. The RT-qPCR reactions were performed on a Stratagene MX3005 using the following parameters: 95°C 5 min; 50 cycles of 95°C 30 sec, 57°C 30 sec, 72°C 30 sec; 72°C 5 min. Fluorescence readings were taken at the end of each elongation step. A melting step was performed following the cycles at 95°C for 30 sec, 54°C for 30 sec and ramp up to 95°C to produce a dissociation curve.

To capture PCR efficiency in the data, Cq values and efficiencies were calculated for each reaction using the LinRegPCR tool (Ramakers et al. 2003, Ruijter et al. 2009). Cq values resulting from efficiencies below 1.80 or above 2.20 were judged unacceptable and were treated as unsuccessful or undetected amplifications. Cq values with efficiency values that were within expected parameters but exceeded (or equaled) 40.00 were also deemed unacceptable and disregarded in downstream analysis. In the same manner, Cq values between (35.00-39.99) were determined to be of low confidence and were marked for special consideration in downstream analysis. Fold-change expression was determined from the Cq values of all gene targets (among all replicates of all samples) among the ‘Bing’, ‘Chelan’, and ‘Skeena’ cultivars using the Pfaffl method (Pfaffl 2001). Expression values were determined with reference to the luciferase spiked gene (Supplementary File 8).

### Short variant identification

The GATK best practices pipeline for short variant discovery was used to identify SNPs and indels in key, differentially expressed ethylene- and auxin-associated contigs, with minor modifications (Van der Auwera et al., 2013; Poplin et al., 2018). Briefly, a group of paired, untrimmed reads from each sweet cherry cultivar was aligned to a designated reference fasta in CLC Genomics Workbench 8.5.1. The reference file contained only the sequences for the assembled contigs that had been previously assigned GO annotations in OmicsBox. The resulting three alignments were exported as BAM files for subsequent use in the GATK pipeline. A reference fasta index and dictionary were created using Samtools and Picard software programs, respectively. Within the GATK (v. 4.1.7.0) suite, the HaplotypeCaller tool was used to identify variants between parental haplotypes for each cultivar; the results from all three cultivars were then merged into a single GVCF file using the CombineGVCFs tool. Finally, The GenotypeGVCFs tool was used to perform joint genotyping on the GVCF file containing variant information for each cherry cultivar. The SelectVariants tool was used to filter out low-quality and low-confidence calls; minimum depth of coverage (DP) was set at 30x and minimum quality of depth (QD), which is directly related to Phred score, was set to 30. The results were visualized and called variants in differentially expressed genes of interest were confirmed, using Integrative Genomics Viewer (v. 2.8.2).

## Results and Discussion

### Pedicel-Fruit Retention Force

Application of ethephon at 80% of fruit development (Smith and Whiting, 2010) ensured developmental equivalency of PFRF and tissue sampling time points across cultivars. This percentage coincided with 12 DBH for ‘Chelan’, 14 DBH ‘Bing’, and 16 DBH for ‘Skeena’. The PFRF values of control fruits decreased naturally over time, with reductions of 75.5%, 74.0%, and 37.5% observed for ‘Skeena’, ‘Bing’, and ‘Chelan,’ respectively. Application of ethephon decreased mean PFRF value in comparison with the respective controls, but whether this decrease was biologically significant and resulted in the achievement of the threshold for mechanical harvesting varied across cultivars (Figure 1).

**Figure 1.**
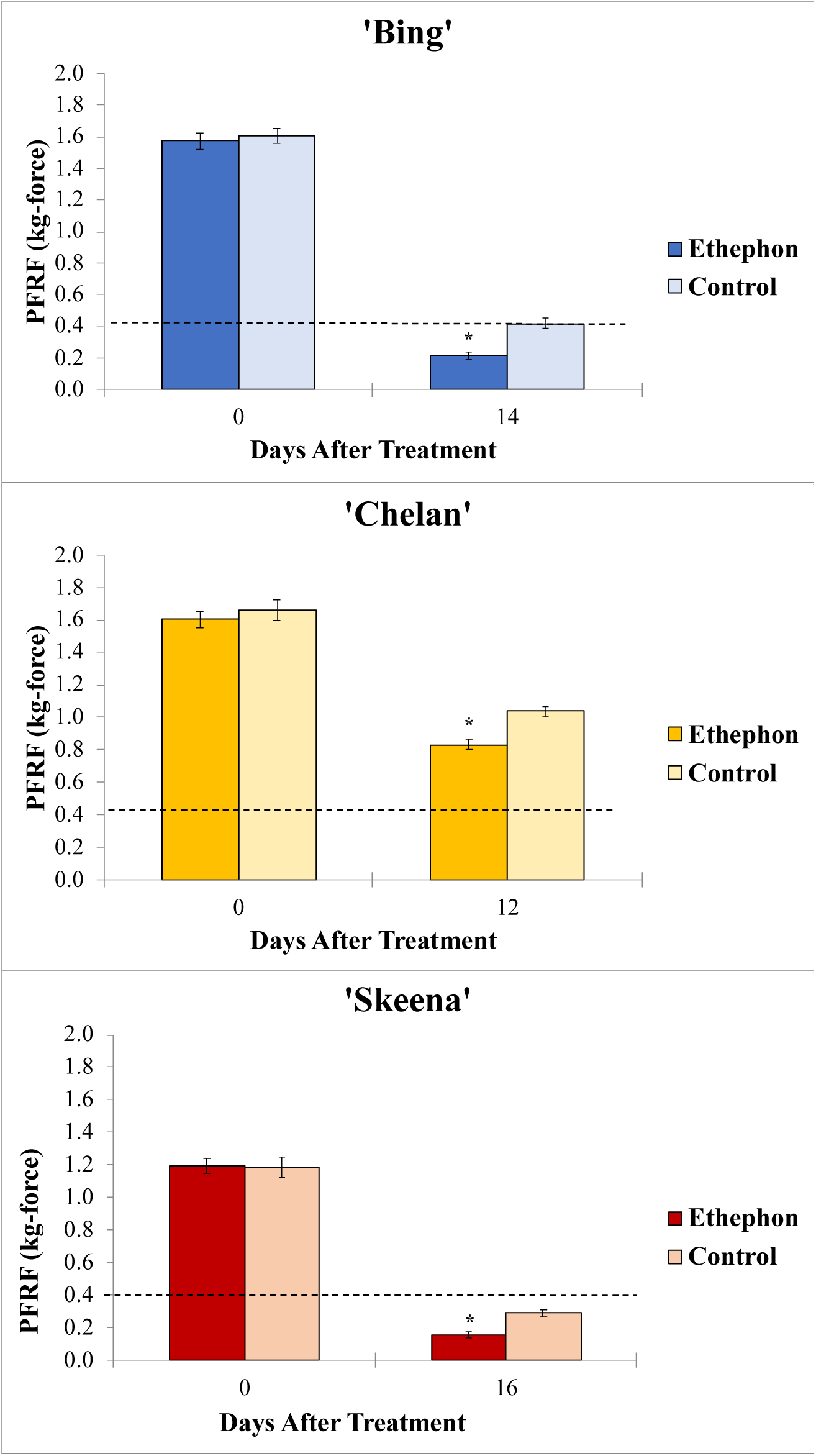
Endpoint mean control and ethephon-treatment PFRF values for (A) ‘Bing’ (B), ‘Chelan’, and (C) ‘Skeena’ PFAZ tissues. The dotted line represents the threshold PFRF for mechanical harvest. Asterisks indicate significant difference between ethephon-treated and control fruit at harvest (p<0.05).

In ‘Skeena’, mean PFRF for both control and treatment fruit dropped below the 0.40kg-force (3.92N) threshold for mechanical harvesting, reaching final mean PFRF values of 0.29kg-force (2.84N) and 0.156kg-force (1.53N), respectively (Figure 1). These findings indicate that while ‘Skeena’ is capable of natural abscission, ethephon application does significantly increase the AZ formation response, causing PFRF values to decrease significantly compared to control fruit by the harvest date.

Similarly, for ‘Bing’, mean PFRF for both control and treatment groups decreased over time; however, a mean PFRF conducive to mechanical harvesting was only achieved in the ethephon treated fruit, which reached a final PFRF value of 0.215kg-force (2.11N), significantly lower than the control value of 0.418kg-force (4.1N) (Figure 1). This suggests that the inducibility of ‘Bing’ is resultant of a similar, yet less dramatic, natural decrease in PFRF that is enhanced by ethephon treatment.

‘Chelan’ exhibited a statistically significant PFRF response at the time of harvest in the ethephon treated fruit in comparison with the control; however, the final PFRF of 0.832kg-force (8.16N) of the treatment group was not reduced to the threshold required for efficient mechanical harvesting (Figure 1). These physiological results support the observations that ‘Chelan’ forms neither a developmental nor an ethylene-induced PFAZ (Smith and Whiting 2010). In the absence of a discrete PFAZ, the pedicel fruit junction region in ‘Chelan’ corresponding to the PFAZ in ‘Bing’ and ‘Skeena’ was used for subsequent RNAseq analysis.

### Transcriptome assembly and annotation

The transcriptome assembly resulted in the generation of 82,587 contigs from 1,061,563,488 total trimmed reads. Contigs were subsequently filtered for >200 base length and >2x coverage, for a final total of 81,852 contigs for downstream processing (Supplementary File 5). Functional annotation conducted using the OmicsBox genomics suite resulted in the assignation of annotations to a total of 30,946 (37.8%) contigs (Supplementary File 7).

### Gene Ontology Enrichment Analysis of Differentially Expressed Contigs

Based on our NOIseq/Kal’s test approach, 1,274 genes were identified to be differentially expressed in ethephon-treated ‘Bing’, 715 in ‘Skeena’, and 523 in ‘Chelan’ at least one treatment/timepoint in comparison with the control. To understand what biological processes, molecular functions, and cellular components were enriched across the three cultivars, and thereby determine pathways of the greatest interest to this study, GO enrichment analysis of differentially expressed contigs was conducted, followed by filtering for the most specific ontologies using an FDR-corrected p-value cutoff of 0.05. This strategy resulted in identification of enriched GOs at 6-hours post-ethephon and at harvest for all genotypes. No GOs were significantly enriched at the 0-hour time point.

### Unique, Enriched Gene Ontologies at 6 Hours Post-Ethephon Treatment

From samples collected 6 hours after treatment, 84 unique GOs were identified for ‘Bing’, 8 for ‘Chelan’, and 33 for ‘Skeena’ (Figures 2A and 3A). Among the ontologies unique to ‘Bing’ were several terms associated with chitin breakdown and metabolism, including: “chitinase activity”, “chitin binding”, “chitin catabolic process”, and “response to chitin”. Besides the hallmark role of plant chitinases and associated pathways in breakdown of fungal pathogens, these enzymes are involved in a number of other important aspects of development, such as generation of signal molecules that regulate organ morphogenesis (van Loon et al., 2006), regulation of lignin deposition and cell shape (Zhong et al., 2002; Johnston, 2006; Agustí et al., 2008), regulation of programmed cell death (Grover, 2012), and formation of abscission zones (Roberts et al., 2000; Grover, 2012). Chitin-associated processes have also been shown to be activated in response to jasmonic acid, methyl jasmonate, ethylene, and gibberellic acid (Graham and Sticklen, 1994; Kasprzewska, 2003; van Loon et al., 2006), as well as auxin and cytokinin (Shinshi et al., 1987). The unique enrichment of chitin-associated GO terms in ‘Bing’ was accompanied by simultaneous overrepresentation of several hormone signaling pathways, including “methyl jasmonate esterase activity”, “methyl salicylate esterase activity”, “jasmonic acid catabolic process”, “salicylic acid metabolic process”, “gibberellin catabolic process”, and “hormone-mediated signaling process”, the latter of which included genes associated with ethylene, auxin, and abscisic acid (ABA) signaling and response. As jasmonic acid (JA) and salicylic acid (SA) metabolic pathways have established roles in plant growth regulation and stress response, the unique co-enrichment of all of these terms, in conjunction with the “ROS response” enriched term, suggest that the inducible abscission phenotype may result from the unique jasmonic acid and redox/ROS signaling pathways, which are known to be activated in response to ethylene response (Müller and Munné-Bosch, 2015). These pathways, in turn, may lead to activation of chitin-associated processes that lead to changes in lignin deposition, cell growth, and programmed cell death in a cascade of events leading up to abscission zone formation following ethephon application in ‘Bing’.

**Figure 2.**
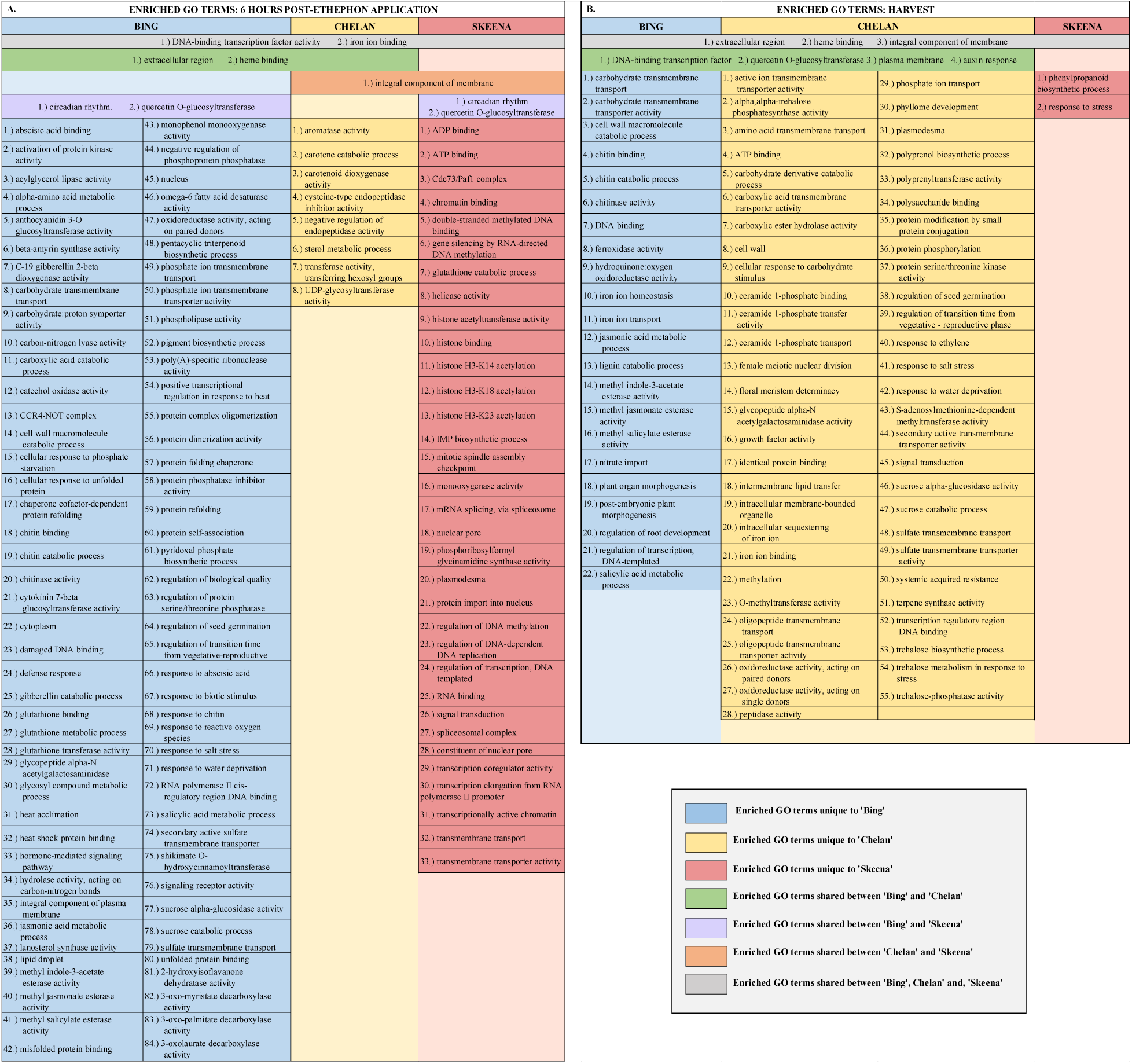
Chart displaying shared and unique enriched GO terms in ethephon-treated ‘Bing’, ‘Chelan’, and ‘Skeena’ at 6 hours post-ethephon treatment (A) and at harvest (B). Enrichment results are based on Fisher’s Exact test with an FDR corrected p-value <0.05.

**Figure 3.**
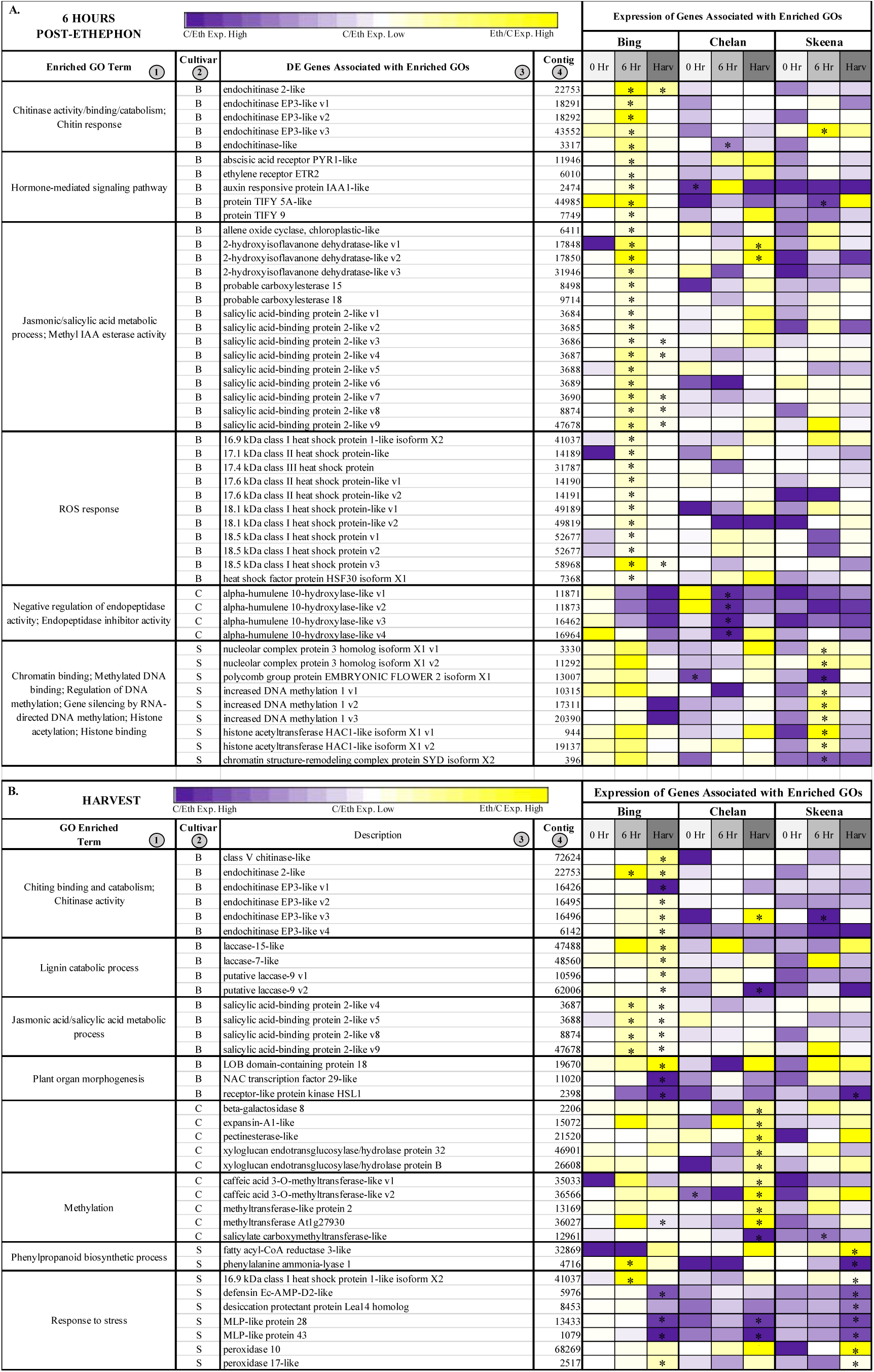
Heatmap displaying expression of genes associated with significantly enriched unique gene ontologies (GOs) for each genotype at 6 hours post-ethephon treatment (A) and at harvest (B). Columns 1 and 2 display enriched GO terms and the genotype for which they were enriched, respectively. Columns 3 and 4 display the names of the differentially expressed genes associated with each enriched GO and their corresponding contig numbers, respectively. Heatmap represents fold change expression of ethephon treated abscission zones (AZs) versus the control AZs for each cultivar. Asterisk indicates significant differential expression (Kal’s Z-test FDR-corrected p-value <0.05, NOISeq DE probability>0.8, |LOG_2_FC|>1).

In ‘Chelan’ a relatively small number of GO terms were enriched at 6 hours post-ethephon. Among the few enriched terms was “negative regulation of endopeptidase activity”. As with plant chitinases, endopeptidases contribute to programmed cell death and cell wall breakdown (Helm et al., 2008) and have been shown to be activated by exogenous ethylene application (LIU et al., 2005). Enrichment of this ontology suggests that molecular mechanisms are at play to stabilize the ethylene response in ‘Chelan’, whereas such a mechanism does not appear to play a substantial role in ‘Bing’.

‘Skeena’ displayed a number of enriched ontologies associated with epigenetic modifications, including “chromatin binding”, “double-stranded methylated DNA binding”, “regulation of DNA methylation”, “gene silencing by RNA-directed DNA methylation”, “histone acetylation”, and “histone binding”. Such processes have previously been implicated in abscission of citrus (Sabbione et al., 2019) and litchi (Peng et al., 2017). In general, methylation of cell wall structural components, such as pectin, is thought to be critical for maintenance of cell wall integrity (Zhang and Zhang, 2020). Enrichment for such processes, which may occur in an ethylene-independent manner, could underlie the natural abscission process in ‘Skeena’.

### Unique, Enriched GOs at Harvest

At harvest, 22 unique enriched ontologies were identified in ‘Bing’, 56 in ‘Chelan’, and 2 in ‘Skeena’ (Figures 2B and 3B). Among the ontologies unique to ‘Bing’ were “lignin catabolic process” and “plant organ morphogenesis”, along with several GO terms that were also enriched at the 6-hour time point— chitin-metabolism and response-associated terms and JA/SA-associated processes. Lignin, a derivative of phenylpropanoid metabolism, is abundant in abscission zones, aiding in the development of a protective boundary layer (Sexton and Roberts, 1982), serving as a mechanical brace to localize cell wall breakdown (Lee et al., 2018), and accumulating specifically in the PFAZ regions during ethylene-promoted abscission (Merelo et al., 2017). In addition to the instrumental role of lignin, general processes involved in “plant organ morphogenesis” appear to be at work in inducible abscission of ‘Bing’.

In ‘Chelan’, a comparatively high number of terms was observed at harvest. Among the enriched GOs were “cell wall-related processes” and “methylation” (Figure 2B, Supplementary File 9). These abscission-associated terms are enriched much later than they are for ‘Bing’ and ‘Skeena’, both of which showed enrichment of associated processes at the 6-hour time point. Furthermore, in contrast to the 6-hour time point, at which few GOs were enriched, the elevated number of enriched terms observed in ‘Chelan’ at harvest suggest a delayed developmental response to ethylene, which may contribute to the failure of this cultivar to develop a PFAZ naturally and in the presence of ethephon.

In ‘Skeena’, at harvest, only two GO terms were uniquely enriched: “phenylpropanoid biosynthetic process” and “stress response”. The limited number of unique terms at the later time point indicate that the majority of impactful processes contributing to PFAZ formation likely take place earlier in development. Among the enriched ontologies, phenylpropanoid metabolism is involved in production of lignin precursors and has been previously implicated in fruit abscission (Kostenyuk et al., 2002). Additionally, stress-associated processes appear to be active at this stage of development and are likely to further accentuate processes associated with programmed cell death, changes in cell wall integrity, and ultimately PFAZ formation in ‘Skeena’.

### Shared, Enriched Gene Ontologies

In addition to unique GOs, shared ontologies reveal PFAZ formation-associated processes that are similar between two or more cultivars following ethephon treatment (Figures 2A, 2B, and 4). At harvest, ‘Bing’ and ‘Chelan’ shared the enriched GO term “response to auxin”, while ‘Skeena’ did not. One possible contributing factor to the auto-abscission of ‘Skeena’ is that this cultivar may have naturally lower free auxin levels than ‘Bing’ or ‘Chelan’ and, promoting natural formation of the PFAZ. This concept is explored further in a later section.

At the 6-hour time point, all three genotypes shared the enriched GO term, “DNA-binding transcription factor activity”. This GO term, among which a number of ethylene-responsive transcription factors were identified (Figure 4), remained enriched in ‘Bing’ and ‘Chelan’ at harvest. This observation provides evidence that all three cultivars have ethylene-responsive capacity; however, the previously described negative regulation of ethylene-associated mechanisms in ‘Chelan’, along with the enrichment of auxin response at harvest, suggests that a higher degree of feedback inhibition serves to quell the extent of ethylene-signaling and response that culminates in strong PFAZ formation in the other two cultivars (naturally, in ‘Skeena,’ and following ethephon stimulation, in ‘Bing’).

**Figure 4.**
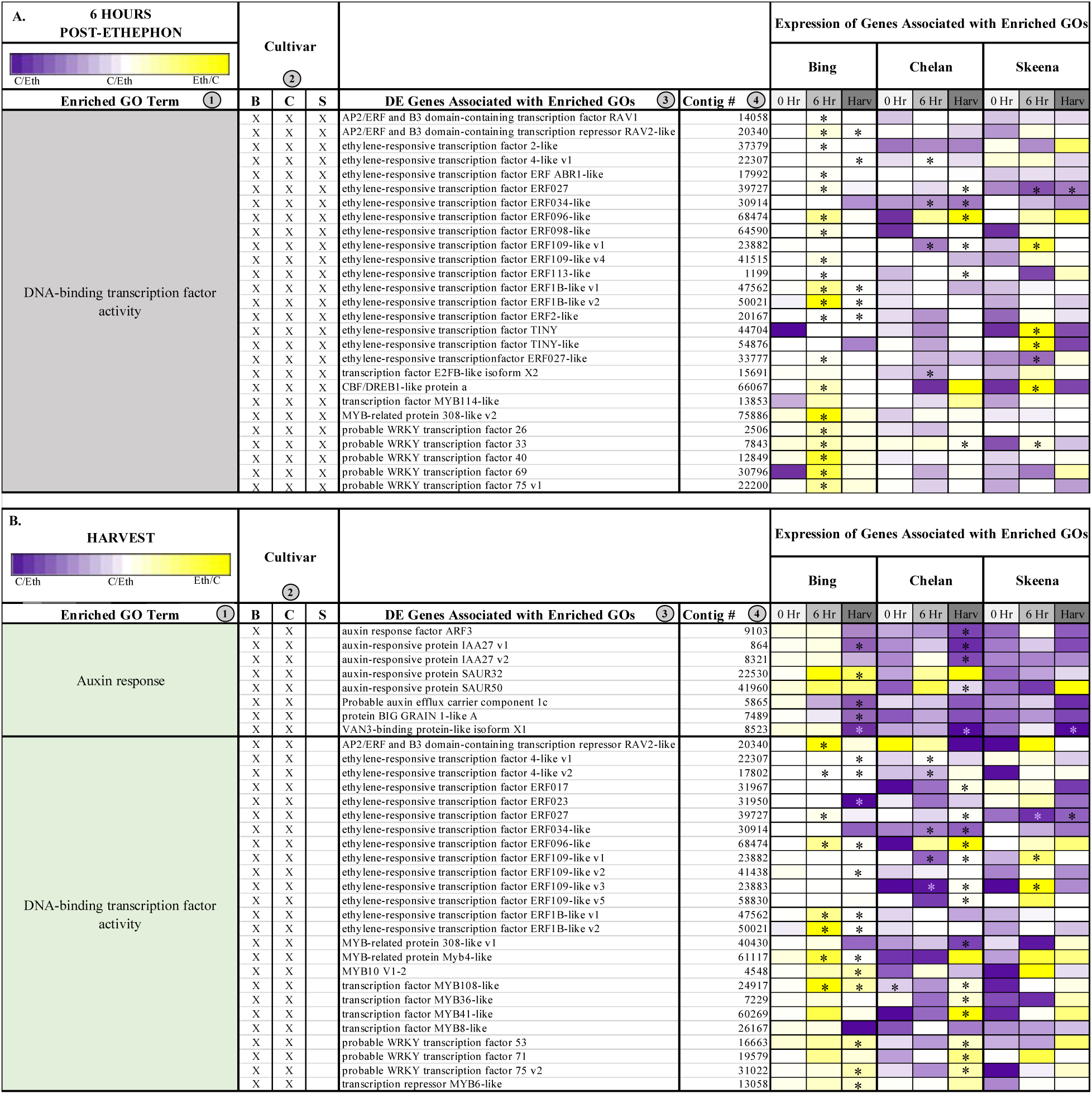
Heatmap displaying expression of genes associated with significantly enriched gene ontologies (GOs) shared between two cultivars 6 hours post-ethephon treatment (A) and at harvest (B). Columns 1 and 2 display enriched GO terms and the genotype for which they were enriched, respectively. Columns 3 and 4 display the names of the differentially expressed genes associated with each enriched GO and their corresponding contig numbers, respectively. Heatmap represents fold change expression of ethephon treated abscission zones (AZs) versus the control AZs for each cultivar.

The term “integral component of membrane” was shared between ‘Chelan’ and ‘Skeena’ at 6 hours post-ethephon and between all three cultivars at harvest. A large number of differentially expressed genes were associated with this term (Supplementary File 9); among the notable genes were cell wall and integrity-associated *patatin-like protein 2* (*PLP2*), which plays a role in programmed cell death, and protein *walls-are-Thin 1* (*WAT1*), which, in addition to its role in auxin mobilization, plays a role in secondary cell wall formation and stability (Camera et al., 2009; Ranocha et al., 2010). As all three cultivars undergo a decrease in PFRF following application of ethephon, it is logical that biological processes, molecular functions, and cellular components associated with membrane and cell wall integrity are affected (Figure 2).

Overall, the GO enrichment results provide a global picture of the processes underlying the natural abscission capacity of ‘Skeena’, the inducibility-of the abscission process in ‘Bing’, and recalcitrance to PFAZ formation in ‘Chelan’. Examination of the differentially expressed genes associated with the enriched ontologies, described in the next section, lends further insight into the mechanisms underlying these three, unique sweet cherry abscission phenotypes.

### Differential Expression of Genes Associated with Key, Enriched Ontologies

Results of the GO enrichment analysis provided a basis for distinguishing pathways in which to observe expression patterns of significant genes, and to compare these patterns of across cultivars following ethephon or control treatments.

### Differential Expression of Genes at 6 Hours Post-Ethephon Treatment

In ‘Bing’, a number of differentially expressed contigs corresponding to endochitinases, including several transcript variants of *endochitinase EP3-like*, *endochitinase-like*, and *endochitinase 2-like* were induced at the 6-hour time point (Figure 3A). At the same time, hormone-mediated signaling genes showed heightened expression, including *abscisic acid receptor PYR1-like* (*PYR1-like*), *ethylene receptor 2* (*ETR2*), *auxin-responsive protein IAA1-like* (*IAA1-like*), and JA responsive protein*s TIFY5A-like* and *TIFY 9*. While, genes associated with chitinase activity, hormone signaling, and ROS response displayed consistently elevated fold-change expression in ‘Bing’, in the other two cultivars, the responses varied. In general, a trend of low or negative fold change expression in ethephon-treated versus control PFAZ tissues was observed for this gene set in ‘Chelan’ and ‘Skeena’; notably, however, *PYR1-like* and *IAA1-like* fold change expression was elevated in both ‘Bing’ and ‘Chelan’. High expression of ABA-associated *PYR1-like,* which promotes senescence and stress responses leading up to abscission, in the PFAZ of ethephon-treated ‘Bing’ and ‘Chelan’ cherries suggests that, despite ultimately different abscission phenotypes, both cultivars possess upstream mechanics that precede PFAZ formation. Furthermore, the low abscission capacity of ‘Chelan’ may lie in a comparatively high auxin versus ethylene response, along with negative regulatory mechanisms that have been previously undescribed. Specifically, several transcripts corresponding to *alpha-humulene 10-hydroxylase-like*, associated with the GO term “negative regulation of endopeptidase activity”, were consistently downregulated in all three cultivars following ethephon treatment, but most drastically so in ‘Chelan’. The specific role of alpha-humulene hydroxylase-like proteins, which modulate plant secondary metabolites (mainly sesquiterpenes) in response to herbivory, in abscission remains to be elucidated; however, the unique expression pattern in ‘Chelan’ points towards a potentially novel mechanism of negative regulation of abscission zone formation.

Many genes associated with epigenetic modification were differentially expressed in ‘Skeena’ at the 6-hour time point, including three transcript variants corresponding to *increased DNA methylation 1* and two transcript variants corresponding to *histone acetyltransferase HAC1-like isoform X1* (*HAC1-like*) (Figure 3A). These genes were also induced in ‘Bing’ at the 6-hour time point, but they were not significantly differentially expressed like in ‘Skeena’. This finding suggests that methylation plays an important role in abscission, and that methylation and associated processes may occur via both ethylene-dependent (‘Bing’) and independent (‘Skeena’) pathways.

### Differential Expression of Genes at Harvest

Among the genes differentially expressed in ‘Bing’ at the harvest time point was “lignin catabolism”-associated *laccase 15-like, laccase 7-like, and laccase 9* (Figure 3B). The former, laccase genes, are involved in lignin synthesis and deposition; these processes are known to occur during AZ formation, and are critical for lignin-promoted architectural changes at the cellular level (Agustí et al., 2008; Merelo et al., 2017). The heightened expression of these genes observed uniquely in ‘Bing’ at harvest suggests that laccases, and therefore structural modifications associated with lignin accumulation, are active in the abscission process for this cultivar.

Also differentially expressed at harvest in ‘Bing’ was “plant organ morphogenesis”-associated *lateral boundary domain containing protein 18* (*LOB18*) (Figure 3B). *LOB18* has previously been implicated in boundary formation during root development. It is possible that this gene, or additional members of the LOB gene family, plays a similar role in boundary formation at the PFAZ (Majer and Hochholdinger, 2011; Wang et al., 2013). *LOB8* expression was high at harvest in both ‘Bing’ and ‘Skeena’, but the expression difference between ethephon-treated and control PFAZ tissues was significant only in the former. In addition to *LOB18*, a differentially expressed transcript corresponding to receptor-like protein kinase *HAESA-LIKE 1* (*HSL1*), which is also associated with the “plant organ morphogenesis” ontology (Figure 3B), was significantly downregulated in both ‘Bing’ and ‘Skeena’. This is consistent with previous work in Arabidopsis that revealed decreased expression levels of HSL1 immediately prior to abscission; furthermore, loss of HSL1 function resulted in impeded floral and leaf abscission (Niederhuth et al., 2013). The differential expression of both *LOB18* and *HSL1* may underlie the ability to induce abscission in ‘Bing’ through boundary formation and promotion of abscission-associated processes.

In addition to the genes that are directly involved in modification of the PFAZ ultrastructure, elevated expression of several transcript variants of corresponding to *salicylic acid binding protein 2-like*, which is associated with JA and SA metabolism, was observed at the harvest time point in ‘Bing’, indicating that JA signaling mechanisms remain active throughout development in ‘Bing’ following ethephon treatment. Similarly, expression of JA-induced, chitinase-associated genes, were also observed at the harvest.

In ‘Chelan’, several genes associated with cell wall modification, including *beta-galactosidase 8*, *pectinesterase-like,* and two *expansin* genes, displayed significantly heightened expression at harvest. These genes are involved in cell wall loosening, degradation of linkages between structural components, and digestion of cell walls and middle lamella in the AZ (Brown, 1997; Cho and Cosgrove, 2000; Roberts et al., 2002; Wu and Burns, 2004). While cell wall modification genes displayed similar expression patterns in ‘Bing’ and ‘Skeena’, the extent of expression change between treatment and control was not significant for these cultivars. Expression patterns of several other genes in ‘Chelan’ indicate that this cultivar may have reduced lignin biosynthetic capacity in comparison to ‘Bing’ and ‘Skeena’. While the expression of cell wall modifying associated genes suggests that ‘Chelan’ is partially competent to develop an PFAZ, it is possible that this cultivar doesn’t possess some of the key molecular equipment to coordinate all of the associated processes, including lignification. This is evidenced by the observation of significantly decreased expression in a methylation-associated transcript corresponding to *caffeic acid 3-O-methyltransferase-like* (*CAOMT*) at harvest; conversely, this transcript displayed increased abundance in both ‘Bing’ and ‘Skeena’ at 6 hours and at harvest. *CAOMT* lies upstream of the lignin biosynthetic process; decreased expression of this gene has been shown to lead to reduced lignification in poplar and expression changes in *CAOMT-like* genes parallels PFAZ formation in citrus (Van Doorsselaere et al., 1995; Jouanin et al., 2000; Merelo et al., 2017).

The expression patterns of genes corresponding to cell wall modifiers and lignin biosynthesis precursors that were identified in ‘Chelan’ lend insight regarding why this non-abscising cultivar displays reduced PFRF but fails to form a PFAZ that is discrete enough to facilitate mechanical harvesting; furthermore, low expression of other terminal abscission-associated genes, like the laccases displaying heightened expression in ‘Bing’ lend further credibility to this hypothesis.

Most notable among the differentially expressed genes in ‘Skeena’, were stress-response associated *peroxidase 10*, *peroxidase 17-like*, and *16.9 kDa class I heat shock protein 1-like.* Elevated expression of stress-responsive genes indicates an increased need for ROS scavenging to prevent oxidative damage while these signal molecules serve the purpose of activating abscission associated processes. Based on the pronounced decrease of stress-responsive genes in ‘Skeena’ following ethephon application, and the observation that PFRF of ‘Skeena’ is reduced the least out of all the cultivars following treatment, it is plausible that ‘Skeena’ may only perceive the exogenous ethylene as environmental stress, whereas ‘Bing’ and ‘Chelan’ appear to host concerted, ethylene-induced signal transduction responses upstream of abscission in response to ethephon application. In conjunction with increased peroxidase gene expression reduced expression of pathogen defense-associated transcripts corresponding to *major latex protein* (*MLP-like protein*) *28* and *43* was observed. This is consistent with a study of *MLP-like* expression patterns in peach abscission zones, wherein it was suggested that *MLP-like* genes play role in mitigation in defense in early AZ formation, but display reduced expression immediately prior to abscission and are further downregulated in response to ethylene (Ruperti et al., 2002). Similar responses were observed for additional plant-defense associated genes, including *dessication protectant protein Lea14*, *defensin Ec-AMP-D2-like*, in both ‘Skeena’ and ‘Bing’.

### Differential Expression of Ethylene-associated Contigs

In addition to observing the expression patterns of differentially expressed contigs associated with key enriched metabolic pathways for each cultivar, it was of interest to specifically examine expression of genes involved in ethylene response, to better understand how these responses may vary at the cultivar level, and thereby differentially contribute to abscission phenotypes. The basis for the inducibility of PFAZ formation in ‘Bing’ during fruit development is further evident at the genetic level with respect to expression of ethylene biosynthesis, signaling, and responsive elements in the transcriptome (Supplementary File 9). Throughout the time course in ‘Bing’, a transcript corresponding to the rate-limiting ethylene biosynthetic gene *1-aminocyclopropane-1-carboxylate oxidase 1 (ACO1*) was highly elevated in expression, as were transcripts corresponding to *ethylene receptor 2* (*ETR2*) and *ethylene insensitive 2* (*EIN2*), two key genes involved in ethylene perception and signaling, respectively. The activation of ethylene biosynthesis that is observed to occur uniquely in ‘Bing’ suggests that exogenous ethephon stimulates the autocatalytic production of endogenous ethylene, which thereby accentuates the overall degree and impact of the ethylene response in this cultivar. Moreover, the naturally high expression of *ETR2* and *EIN2*, which are augmented in the presence of exogenous ethephon lend further testament to the responsiveness of ‘Bing’ to ethylene. A negative regulator of ethylene, *reversion-to-ethylene-sensitivity 1* (*RTE1*) was induced in ‘Bing’ following ethephon application, but levels remained lower overall than in ‘Chelan’ and ‘Skeena’, an observation which is consistent with the comparatively high degree of sensitivity of ‘Bing’ to ethylene. In contrast to ‘Bing’, ‘Skeena’ and ‘Chelan’ displayed basal levels of expression of *ACO1* and *ETR2*, and elevated levels of *RTE1*, suggesting minimal ethylene response, endogenous production, and sensitivity (Figure 5, Supplementary File 9).

**Figure 5.**
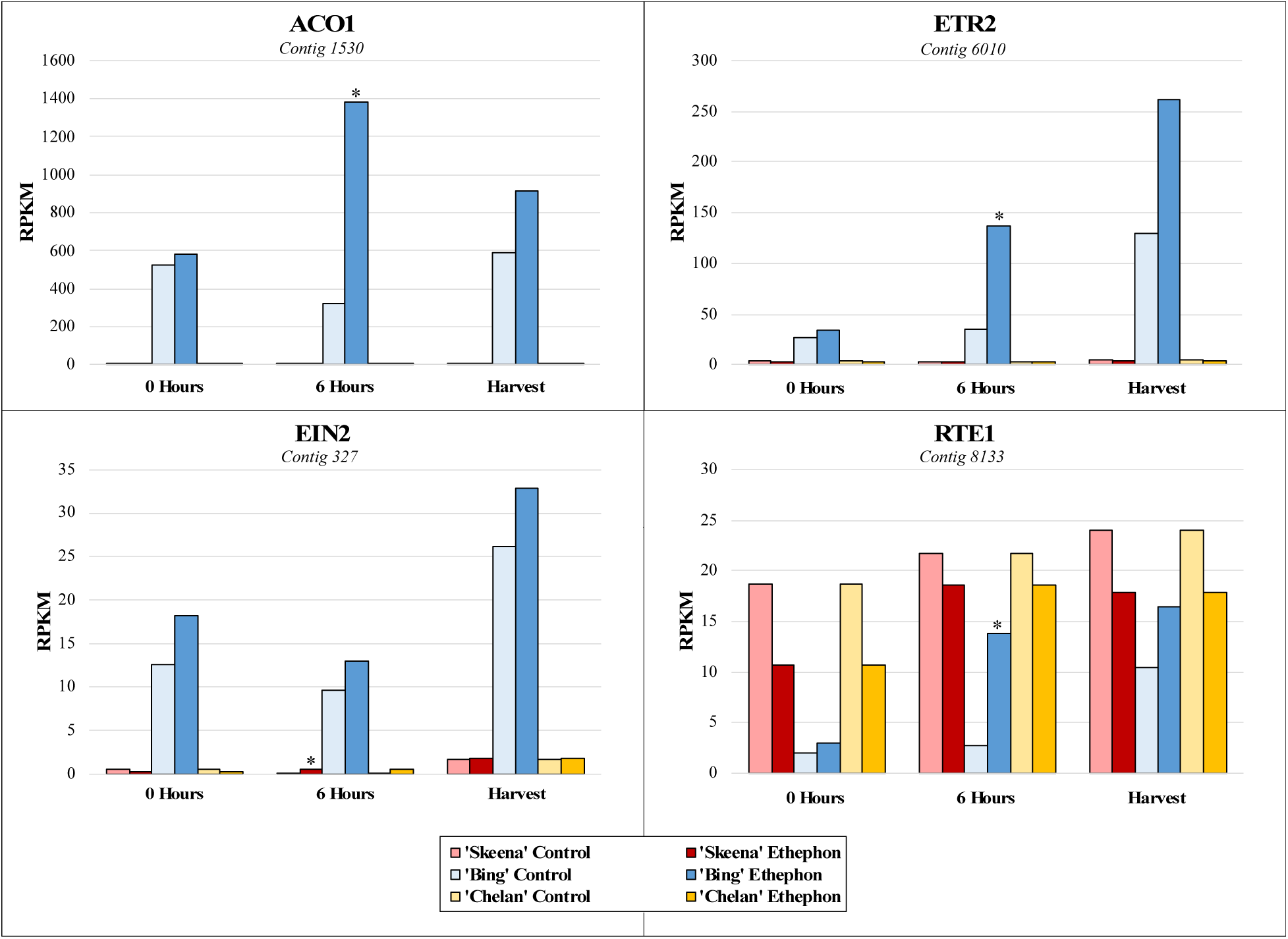
Expression patterns of ethylene biosynthesis gene *1-aminocyclopropane-1-carboxylate oxidase 1* (*ACO1*), *ethylene receptor 2* (*ETR2*), and signal transducer *ethylene insensitive 2* (*EIN2*), and ethylene negative regulatory protein *reversion-to-ethylene-sensitivity 1* (*RTE1*) at 0 hours, 6 hours post-ethephon treatment, and harvest. Asterisks indicate significant difference, according to both Kal’s Z-test (p<0.05) and NOIseq-sim (probability > 0.8) differential expression probability analysis, and a |logFC|>1.

Transmission of ethylene-activated signal to the nucleus involves ethylene-insensitive 2 (EIN2) and ethylene insensitive 2-like (EIL) family of proteins. Upon activation of ETRs by ethylene, the C-terminal domain of EIN2 is cleaved and translocated to the nucleus where it activates transcription of ethylene-responsive factors (ERFs) involved in the regulation of downstream ripening responses (Liu et al., 2015; Chen et al., 2018). Numerous transcripts encoding ERFs were identified, which were differentially expressed in at least one cultivar and time point (Figure 4). Consistent with the aforementioned results, a majority of these contigs were highly induced in ‘Bing’ following ethephon treatment. In particular, transcripts corresponding to *ERF1B-like*, *ERF27* and *ERF109-like, ERF113-like*, *ERF109-like,* and *ERF ABR1-like* were highly differentially expressed in ‘Bing’ 6 hours following ethephon treatment, and also at harvest. In all three cultivars *ERF96-like* was highly induced following ethephon treatment (Figure 4, Supplementary File 9). In addition to ERFs, transcripts corresponding to members of the of the WRKY and MYB transcription factor families displayed highly elevated expression in ‘Bing’ following ethephon treatment. This is consistent with previous work demonstrating the activation of both of these transcription factor families during abscission, and eliciting actions downstream of initial ethylene response (Corbacho et al., 2013; Li et al., 2015; Jiang et al., 2017; Zhang et al., 2019).

### Differential Expression of Auxin Response-associated Contigs

In addition to the differential ethylene responses, the ethephon-treated ‘Chelan’, ‘Bing’, and ‘Skeena’ PFAZ tissues displayed a differential reduction in the expression of auxin-responsive genes in comparison to their respective controls. This suggests that the response to auxin and/or reduction in auxin transport, was inhibited in the presence of exogenously applied ethylene.

Auxin-responsive transcription factor genes *IAA11*, *IAA13*, and *IAA27*, auxin transport facilitator *WAT1*, and auxin efflux carrier component 1c displayed comparatively low expression in ‘Skeena’, intermediate expression in ‘Bing’, and higher expression in ‘Chelan’ control PFAZ tissues at 6 hours post-ethephon and at harvest. Ethephon treatment attenuated this expression in PFAZ tissue for all three cultivars, but the same relative expression trend was maintained. *IAA13*, *IAA27*, and *WAT1* displayed decreased expression at harvest in all ethephon-treated cultivars in comparison with their respective controls. Interestingly, ‘Chelan’ displayed a natural increase in the expression of these transcripts from the 0-hour time point to harvest, which was inhibited and reversed by the application of ethephon (Figure 6). For both ‘Bing’ and ‘Skeena’, the same transcripts naturally decreased in abundance over time—a decrease that was accelerated in the presence of ethephon.

**Figure 6.**
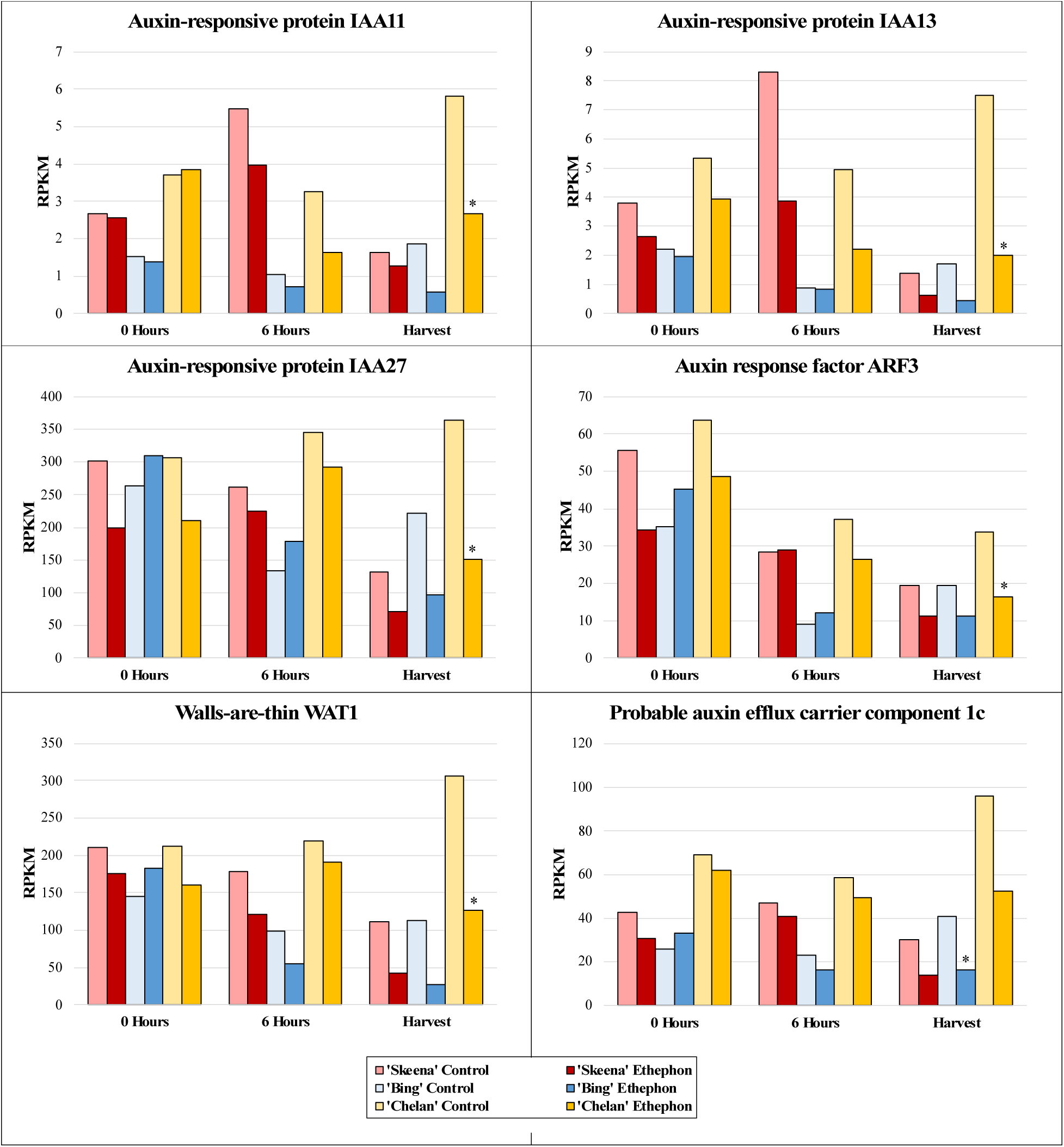
Expression patterns of auxin-associated genes. A shared expression pattern was observed for auxin-associated genes at harvest. Auxin responsive transcription factors *IAA11*, *IAA13*, and *IAA27*, auxin transport facilitator *walls-are-thin 1* (*WAT1*), and auxin efflux carrier component 1c displayed comparatively lower expression in ‘Skeena’, intermediate expression in ‘Bing’, and higher expression in ‘Chelan’ control at harvest. Following ethephon treatment, expression was attenuated in all three cultivars, however the same relative expression trend was maintained, with ‘Skeena’ ethephon-treated fruit exhibiting lowest expression at harvest and ‘Chelan’ ethephon-treated fruit displaying the highest expression at harvest. Asterisks indicate significant difference, according to both Kal’s Z-test (p<0.05) and NOIseq-sim differential expression probability analysis (probability>0.8), and a |logFC|>1.

While free auxin was not measured in this study, differences in the abundance of auxin-responsive and auxin mobilization-associated transcripts in the three sweet cherry cultivars over time lends to extrapolation of information regarding cultivar-specific, endogenous free IAA concentrations. If free auxin levels are high, abundance and/or activity of auxin-responsive protein encoding transcripts (ARFs, IAAs) is expected to be higher to accommodate them. ‘Skeena’ does not require ethylene for abscission; however, ethephon application appeared to further offset the auxin to ethylene ratios in favor of ethylene to accelerate abscission in this cultivar. The ability of ‘Skeena’ to auto-abscise suggests that endogenous accumulation of free auxin at the site of abscission may be naturally lower in this cultivar than it is for ‘Bing’ and ‘Chelan’. Conversely, the comparatively higher expression of transcripts associated with auxin response and movement proteins in ‘Chelan’ is suggestive of a naturally higher level of endogenous IAA. While the current industrial standard levels of ethephon application were insufficient to reduce the PFRF of ‘Chelan’ fruit to the threshold required for mechanical harvesting, both PFRF and auxin-associated transcript abundance in ‘Chelan’ did decrease as a result of ethephon application. Ethephon application rates as high as 5.8 L ha^−1^ remain insufficient to induce a reduction of ‘Chelan’ PFRF values to the threshold for mechanical harvest (Smith and Whiting, 2010). An early study found that a high application rate of 500 ppm (7.8 L ha^−1^ [6.7 pt A^−1^]), ethephon begins to deleteriously affect some sweet cherry cultivars by inducing unwanted leaf abscission and terminal shoot necrosis, although it is not reported to what extent ‘Chelan’ is impacted by such a concentration (Bukovac and MJ, 1979; Smith and Whiting, 2010). Considering the present results alongside the insight gained from previous work, it is possible that mechanical abscission of ‘Chelan’ could be achieved through a combination of a slightly higher ethephon application rate and application of auxin inhibitors, the latter of which could further shift the ethylene/auxin ratio in a manner favorable to abscission while reducing the need for excessively high and potentially phytotoxic ethephon application rates.

‘Bing’ PFAZ tissues displayed an abundance of auxin-associated transcripts intermediate to that of ‘Skeena’ and ‘Chelan’. Following ethephon treatment, the phenotypic observations of inducible abscission were supported at the gene expression level; with the responses greatly attenuated to lower than those of ‘Skeena’ ethephon treated and control PFAZ tissues.

### Proposed Models for PFAZ Formation

Together, the GO enrichment and differential expression analysis results inform a new model for abscission (Figure 7). In ‘Bing’, ethephon elicits an ethylene-response that stimulates hormonal regulatory signals and metabolic processes, including JA and SA-mediated signaling. These processes, in turn, activate downstream processes associated with morphological changes at the PFAZ, including chitin metabolism and lignin biosynthesis and deposition. The culmination of these events ultimately facilitates mechanical separation of the fruit at the PFAZ. The components of this process, which appears complete in ‘Bing’, seem to be partially lacking in ‘Chelan’, perhaps due to inherently high auxin levels that limit the ethylene-response capacity from the beginning. ‘Skeena’ appears to orchestrate PFAZ formation in via a different and ethylene-independent mechanism. While this mechanism will require further elucidation, lack of enrichment for the “auxin response” ontology and reduced expression of the corresponding auxin-associated genes in ‘Skeena’, suggests that low levels of free auxin, and therefore minimal antagonism of a natural abscission process, may underly the auto-abscission phenotype.

**Figure 7.**
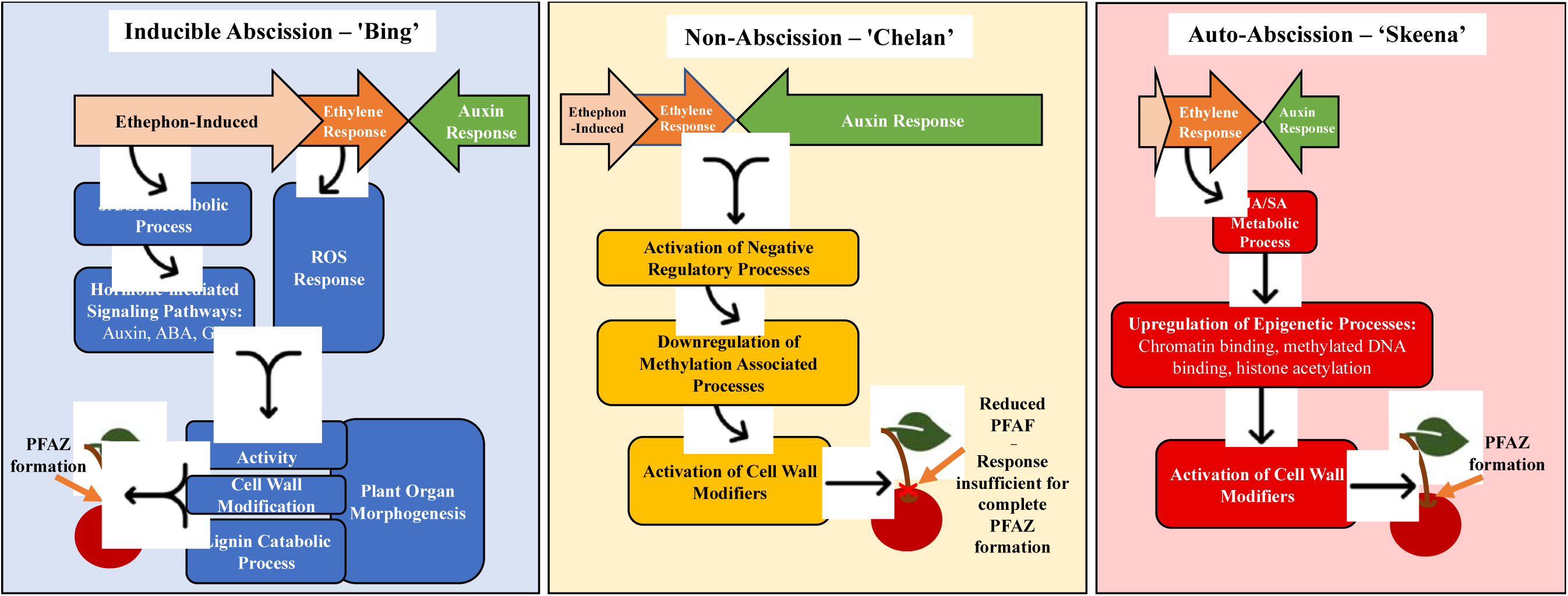
Proposed models for abscission, based on gene expression and gene ontology (GO) enrichment results following ethephon application of ‘Bing’, ‘Chelan’, and ‘Skeena’ sweet cherry. In ‘Bing’ (A), ethephon offsets the balance between ethylene and auxin in favor of ethylene. This triggers endogenous ethylene-responses that stimulates JA and SA-mediated signaling, additional hormone mediated signaling pathways, and reactive oxygen species (ROS) responses. Together, these ethylene-induced signaling pathways activate downstream processes associated with morphological changes at the PFAZ, including chitin metabolism, cell wall modification, plant organ morphogenesis, and lignin biosynthesis and deposition. The culmination of these events ultimately facilitates mechanical separation of the fruit at the PFAZ. In ‘Chelan’ (B), the components required for inducible abscission seem to be partially lacking. While ethephon likely contributes to enhanced ethylene signaling, free auxin levels in ‘Chelan’ may be high enough initially to overwhelm any ethylene response, thus limiting the cascade of signaling events leading to abscission. The activation of cell wall modifying enzymes may be responsible for the observed reduction in PFRF, however, it does not seem to be enough to facilitate complete PFAZ formation. In ‘Skeena’, ethephon does little to affect the overall abscission phenotype, as this cultivar is capable of natural abscission. While ethylene signaling is not high, low initial levels of free auxin may naturally shift the hormonal tug-of-war in favor of abscission, leading to moderate activation of JA metabolism and signaling. Near the time of harvest, upregulation in processes associated with epigenetic modification and, ultimately, activation of cell wall modifiers may represent intermediates in an alternate, ethylene-independent pathway to PFAZ formation.

### Short Variant Discovery in Differentially Expressed Genes

Variant calling using the GATK pipeline revealed several SNPs and indels in the ORFs of DE genes associated with key, enriched GOs (Table 1).

**Table 1.**
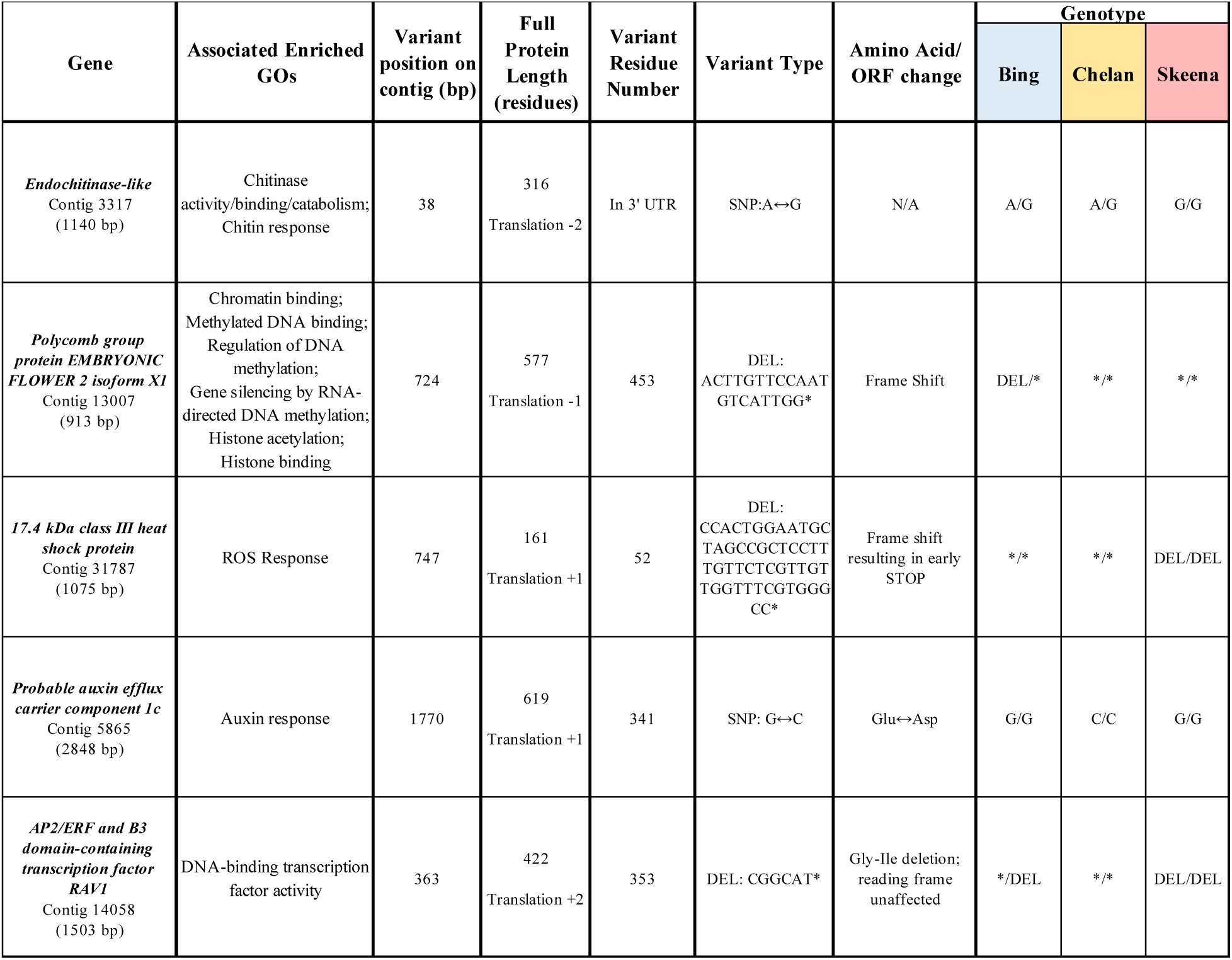
Variants identified in differentially expressed genes associated with enriched gene ontologies. The GATK RNAseq short variant discovery pipeline was used for characterization of SNPs and indels (Poplin et al., 2018).

An INDEL was identified in “DNA-binding transcription factor activity”-associated *AP2/ERF and B3 domain containing protein RAV1*, wherein deletion of a 6 bp segment results in loss of two amino acid residues without impacting the reading frame. Interestingly, ‘Skeena’ was homozygous for this deletion, ‘Bing’ was heterozygous, and ‘Chelan’ did not possess the deletion. The resulting protein changes likely impact the natural propensity of the cultivars to transduce signals associated with stress and hormone responses upstream of PFAZ formation and could ultimately underlie differences in abscission phenotype. Additionally, a SNP, resulting in an amino acid substitution, in the coding sequence of *auxin efflux carrier component 1c*, was identified in ‘Chelan’. This variant could impact the overall auxin responsiveness, which appears to be greater in ‘Chelan’ than ‘Bing’ or ‘Skeena’, at the genetic level.

Additional variants identified include a SNP in “ROS response”-associated *17.4 kDA class III heat shock protein*, an INDEL in the epigenetic modification-associated gene *polycomb group EMBRYONIC FLOWER 2 isoform x1,* and a SNP in the 3’ UTR of the “chitinase activity”-associated *endochitinase-like* gene. *A* member of the diverse polycomb group (PcG) family, *EMF2* is involved in epigenetic silencing of developmental repressors, and was significantly downregulated in ‘Skeena’ at the 6-hour time point. While further work is needed to determine the precise nature of these variants and their impact on PFAZ formation, their presence in DE genes associated with enriched functions following ethephon application makes them key targets for understanding activation of abscission in sweet cherry cultivars.

### RT-qPCR validation

RT-qPCR analysis of 10 ethylene-responsive/abscission-related genes resulted in 70% correspondence of general expression trends for ‘Bing’ and ‘Chelan’ and an 80% correspondence for ‘Skeena’. Validated transcripts whose RT-qPCR expression pattern (fold-change calculated using the 2^-(ΔΔ^ method) was consistent with that of the RNAseq-based expression (ethephon/control RPKM ratio) included genes associated with ethylene biosynthesis (ACS1 and ACO1), perception (ETR2), and response (ERF1B-like, ERF027-like, and WRKY1), as well as cell wall breakdown-associated polygalacturonase (PG) (Supplementary File 8).

### Conclusions

This study investigated the changes in PFRF, gene expression changes, and enriched biological processes underlying PFAZ formation in sweet cherry. The results provide transcriptomic insight regarding the gene expression-level effects of exogenous ethylene application on the abscission phenotype and PFAZ development, which can be utilized by the industry to customize harvest strategies.

Consistent with previous work, PFAZ formation was observed to occur in a cultivar-specific manner, and the abscission phenotype of each cultivar was affected to a different degree by exogenous application of ethylene. Observation of unique, heightened expression of ethylene biosynthesis, perception, signaling, and response genes in ‘Bing’ at 6 hours following ethephon application and at harvest parallels the decrease in PFRF and may be partially responsible for the inducibility of abscission in this cultivar. GO enrichment analysis revealed key biochemical pathways containing additional differentially expressed genes potentially underlying the unique, abscission responses of each cultivar. Among these enriched pathways were a number of hormone-signaling pathways, ROS signaling, chitin metabolic processes, and lignin biosynthesis, and DNA-binding transcription factor activity (among which were numerous ERFs).

Overall, the results of this study point towards a potential genetic basis for the inducible abscission response in ‘Bing’, the auto-abscission of ‘Skeena’, and the recalcitrance to abscise displayed by ‘Chelan’. The natural abscission of ‘Skeena’, which is enhanced by ethephon, may result from naturally lower levels of free auxin, as evidenced by the low abundance of IAA/ARF-associated transcripts in comparison with ‘Bing’ and ‘Chelan’. Furthermore, ‘Skeena’ exhibited fewer significant gene expression changes than the other cultivars following ethephon application, which corresponded to fewer enriched ontologies at the time of harvest. In ‘Chelan’, comparatively high abundance auxin-associated transcripts in control fruit, that was attenuated following ethephon treatment, may indicate higher levels of endogenous free auxin. The increased capacity for auxin mobilization, indicated by IAA/ARF expression, antagonizes the abscission-promoting effects of exogenous ethylene. This observation, along with the PFRF results, provides insight regarding the recalcitrance of ‘Chelan’ fruit to abscise at the pedicel-fruit junction.

Identification of cultivar-specific, differentially expressed genes and enriched GO terms involved in abscission, as well as ontologies that are shared across cultivars, provides information that will inform future efforts to promote controlled, timely abscission of sweet cherries. This, in turn, will lead to the improvement and standardization of mechanical harvesting, thereby improving efficiency and increasing the economic profitability of the sweet cherry industry. Ultimately, the outcomes of this work may be extended to other crops where planned abscission can be useful in managing the harvest.

## Acknowledgments

This study was supported by a USDA Special Crop Research Initiative (SCRI) program grant (Project No. 2009-02559) to MW and AD. Work in the Dhingra lab was supported in part by Washington State University Agriculture Center Research Hatch Grant WNP00011. SLH, BK, and TK acknowledge the support received from ARCS Seattle Chapter and National Institute of Health/National Institute of General Medical Sciences through an institutional training grant award T32-GM008336. The contents of this work are solely the responsibility of the authors and do not necessarily represent the official views of the NIGMS or NIH. The authors thank Dr. Richard Sharpe for assistance with RT-qPCR assays and analysis. This manuscript has been released as a pre-print at BioRxiv (Hewitt, Kilian, et al. 2020).

## Author Contributions

SH – Transcriptome assembly, data analysis, manuscript preparation, and editing

BK – Experimental design, data analysis, manuscript preparation

TK – Experimental design, PFRF, and tissue collection

JA – Assisted with ethephon treatment and collection of PFRF data and tissue collection

MW – Experimental design, guidance on application of ethephon and tissue collection

AD – Conceived the study, experimental design, data analysis, manuscript preparation and editing

## Competing Interest

The authors declare no competing interests, or other interests that might be perceived to influence the results and/or discussion reported in this paper.

**Supplementary File 1.** Ethephon treatment and sampling timeline

**Supplementary File 2**. PFAZ data from 2010, 2013, and 2014 seasons

**Supplementary File 3.** Supplemental fruit color data collected at the same time as PFRF and AZ tissue sampling.

**Supplementary File 4.** Picture of fruit section sampled for PFAZ analysis

**Supplementary File 5.** Cherry assembly fasta file

**Supplementary File 6.** Spreadsheet with RPKM values of contigs that passed stringent statistical and probability filters for differential expression based on Kal’s Z-test, NOIseq-sim analysis, and Log2FC expression filtering.

**Supplementary File 7.** List of all annotated contigs >2x coverage and >200bp in length

**Supplementary File 8.** RT-qPCR primers; Cq values and fold change expression summary by template

**Supplementary File 9.** Expression heatmaps for differentially expressed (DE) genes associated with shared and unique enriched gene ontology (GO) terms and DE genes associated with ethylene and auxin pathways.

