## Supplementary figures and images for "Characterization of ethylene-inducible pedicel-fruit abscission zone formation in non-climacteric sweet cherry (*Prunus avium* L.)"

### Supplemental File 1

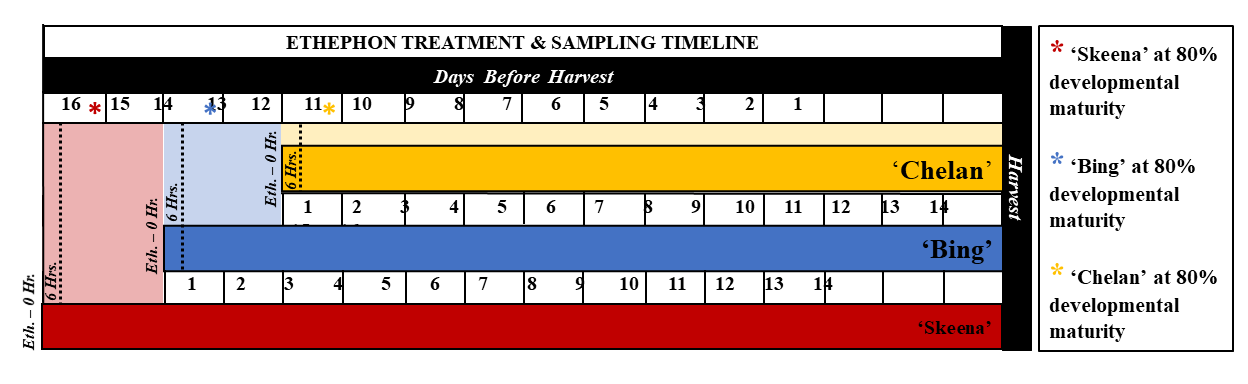

### Supplemental File 4

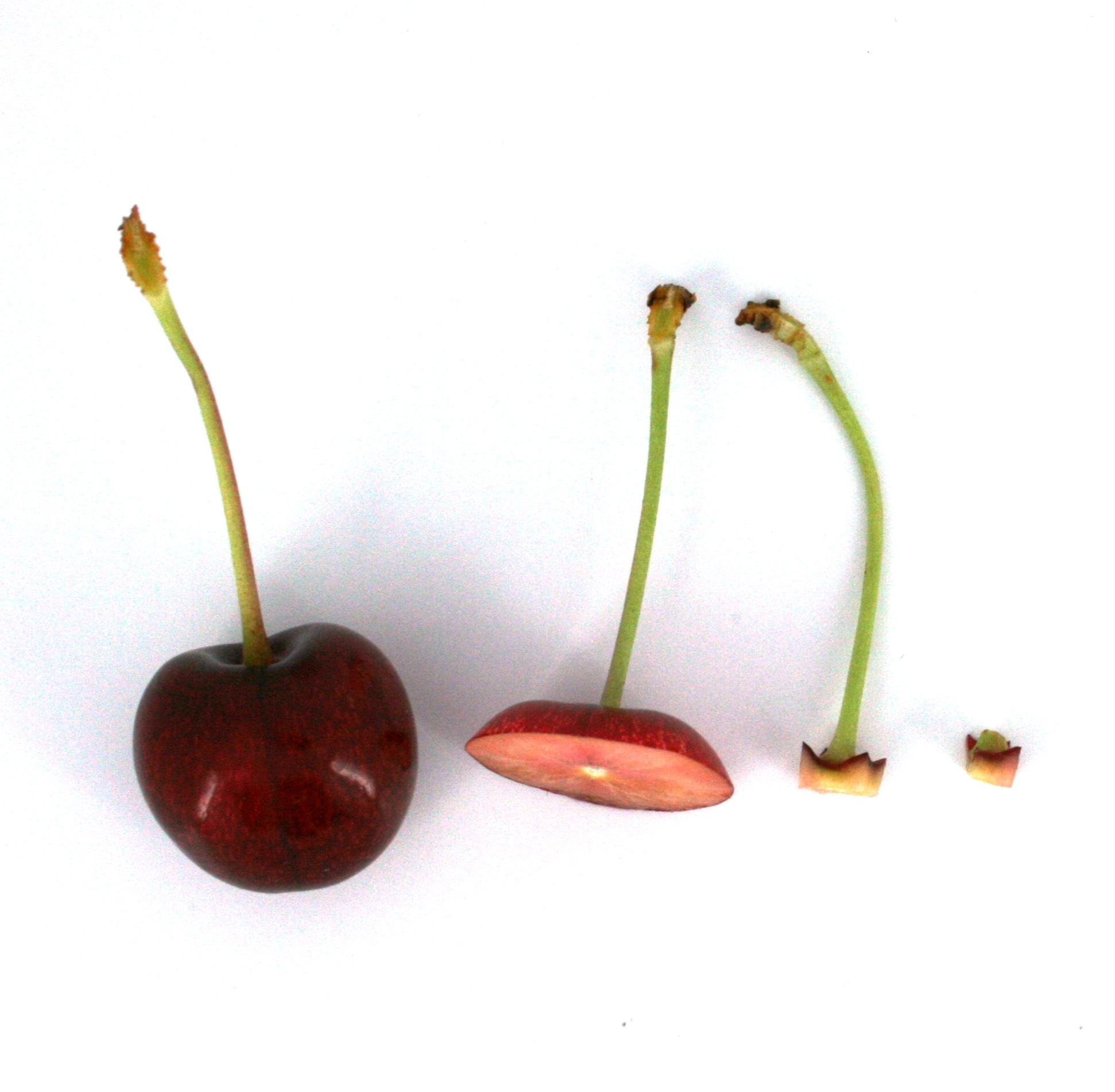
